# A cohesion optimum underlies chromosome segregation fidelity in oocytes

**DOI:** 10.64898/2026.05.08.723885

**Authors:** Yan Yun, Kanako Ikami, Duy Do, Ziyang Guo, Jiyeon Leem, Hilina Bekele, Binyam Mogessie, Neil Hunter

**Affiliations:** Howard Hughes Medical Institute; University of California Davis, Davis, United States; Department of Microbiology & Molecular Genetics, University of California Davis; Davis, United States; Center for Reproductive Medicine, Shantou Clinical Medical College of Jinan University (Shantou Central Hospital), Jinan University, Shantou, Guangdong, China; Department of Molecular, Cellular and Developmental Biology, Yale University, New Haven, CT, USA; Department of Molecular & Cellular Biology, University of California Davis; Davis, United States

## Abstract

Chromosome segregation is compromised in eggs from women of both early and advanced reproductive ages. Deteriorating cohesion causes premature separation of sister-chromatids in eggs from older females. We show that the converse is true for oocytes of adolescents, with excessive cohesion impeding segregation. Oocytes from juvenile mice show severe chromosome lagging in anaphase I, leading to nondisjunction or, in extreme cases, failure of the first meiotic division. These defects are suppressed by experimentally weakening cohesion or enhancing its resolution during anaphase I. By contrast, lagging and nondisjunction are rare in the oocytes of young adults because cohesion is inherently weaker. Thus, relative cohesion strength underlies both the frequency and type of segregation errors observed in eggs throughout the female reproductive lifespan.

**One-Sentence Summary:** In eggs, errors in chromosome segregation arise from age-dependent imbalances in how tightly chromosomes are held together.

---

Oocyte meiosis initiates *in utero* during which homologous chromosomes (homologs) become connected by chiasmata, the conjunction of crossing over between homologs and cohesion between sister chromatids, which is mediated by cohesins–large multi-subunit ring proteins that topologically entrap newly replicated sister chromatids (*1-7*). Oocyte meiosis then arrests around birth, and a finite ovarian reserve of quiescent primordial follicles is established. Following the resumption of meiosis in fully grown follicles, chiasmata enable stable bipolar attachment of homolog pairs to the spindle, which is required for accurate segregation at the first meiotic division (MI). Homolog segregation ensues when separase is triggered to resolve chiasmata by cleaving the cohesin rings that connect chromosome arms (*8*). Centromeric cohesion is protected from cleavage until the second meiotic division (MII) when sister chromatids segregate.

The well-documented decline in egg quality in women of advanced reproductive age, known as the maternal-age effect, is largely attributed to chromosomal errors caused by the premature separation of sister-chromatids due to progressive weakening of cohesion during the protracted arrest of primordial follicles (*9-17*). Hoffmann and colleagues made the striking observation that eggs from adolescent females also have high levels of chromosomal errors, almost exclusively aneuploidy resulting from homolog nondisjunction in which homologs fail to separate at MI (*18*). The high level of aneuploidy reported in oocytes from juvenile pigs (*19*) suggests that this phenomenon may be conserved in mammals and prompted us to analyze chromosome segregation in oocytes from juvenile mice (J-oocytes, Fig.1 and fig. S1). Live imaging of J-oocytes from 3-week-old mice revealed that 92.2% showed chromosome lagging during anaphase I compared to only 15.1% of oocytes from young sexually-mature adults (3-months old; Y-oocytes; Fig. 1, A and B; *p*=0.0022, Mann-Whitney test; fig. S1, A to E; Supplementary Movies 1, 2). Moreover, failure of MI, indicated by polar-body retraction (PBR) occurred in 16.6% of J-oocytes but was never observed in Y-oocytes (fig. S1, B, F and G). Aneuploidy was observed in 23.3% of J-oocytes that completed MI compared to just 0.8% in Y-oocytes (Fig. 1, C and D; *p*<0.0001, Fisher’s exact test). Moreover, aneuploidy was exclusively due to nondisjunction and often involved multiple chromosomes in the same nucleus (fig. S1H). This contrasts chromosomal abnormalities seen in oocytes from mice of advanced maternal age (15-19 months, A-oocytes), which are predominantly pre-division errors that result in the appearance of free chromatids in MII-arrested cells (Fig. 1, C and D). This pattern of high nondisjunction in oocytes from juvenile females and high pre-division errors in oocytes from reproductively advanced females is analogous to that observed in humans (Fig. 1E and fig. S1I; adapted from (*18*), pointing to conserved underlying mechanisms.

**Figure 1.**
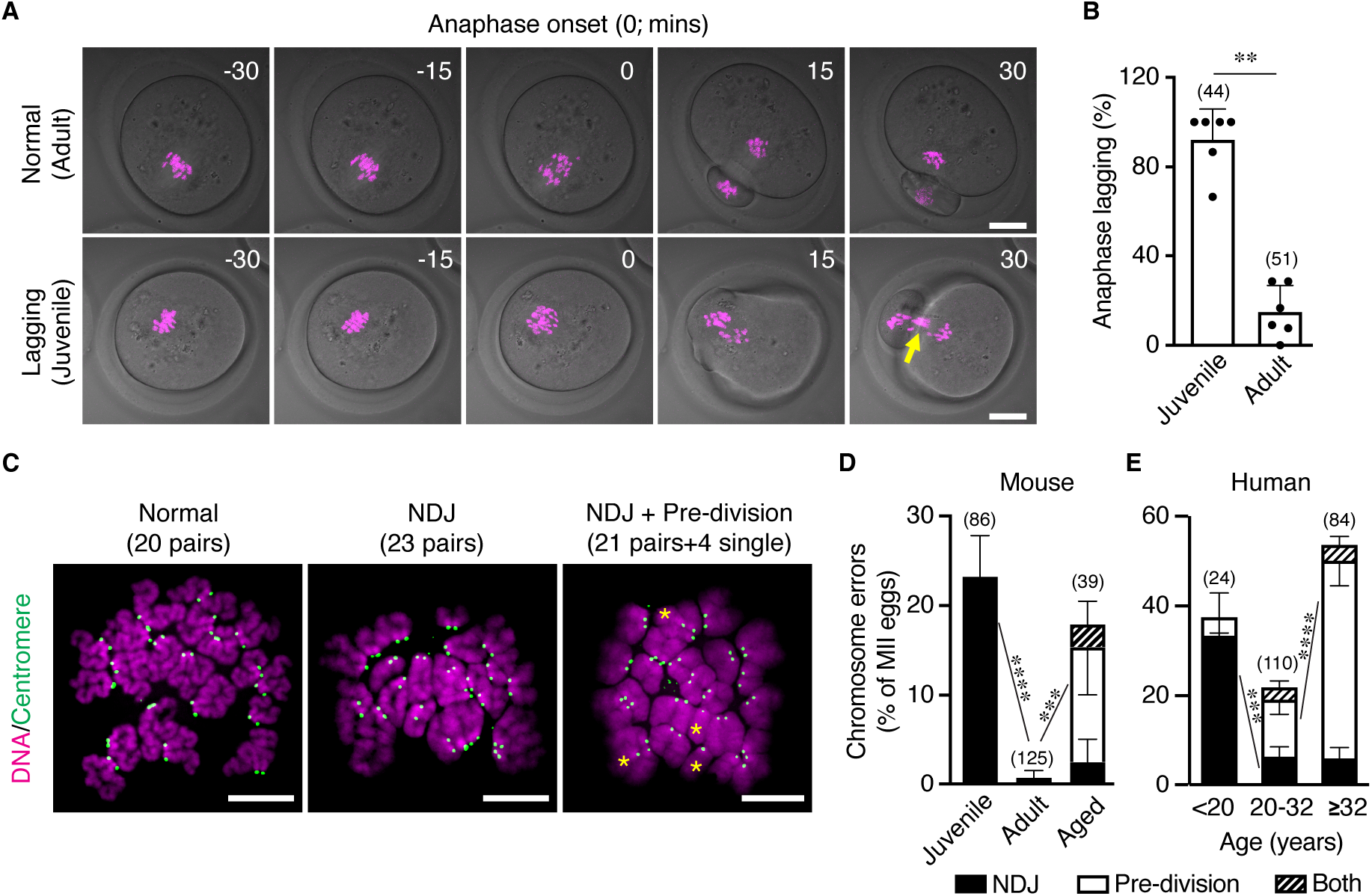
Chromosome lagging and homolog nondisjunction errors in oocytes from juveniles. **(A)** Representative frames from live-cell imaging of a Y-oocyte (from a young adult) and a J-oocytes (from a juvenile) spanning the anaphase I stage. The yellow arrow highlights chromosome lagging. Scale bars, 20 µm **(B)** Quantification of chromosome lagging in anaphase I from the experiments represented in A. Each data point represents an individual experiment. Error bars indicate SD. **(C)** Representative images of metaphase-II (MII) oocyte chromosome spreads. Asterisks highlight single chromatids indicative of pre-division errors. NDJ, nondisjunction. Scale bars, 10 µm **(D)** Frequency and type of chromosome errors in mouse MII eggs from juvenile, young adult, and aged animals. Errors bars show standard error of a proportion. ***, *p* <0.001; ****, *p*<0.0001; Fisher’s exact test **(E)** Frequency and type of chromosome errors in human MII eggs from women in the indicated age groups (adapted from Gruhn et al. {Gruhn, 2019 #3553}). Errors bars show standard error of a proportion. ***, *p* <0.001; ****, *p*<0.0001; Fisher’s exact test

Chromosome lagging and nondisjunction at the first meiotic division imply that connections between homolog are inefficiently resolved in J-oocytes. Therefore, we asked whether weakening cohesion or enhancing its resolution could suppress these defects (Fig. 2). Heterozygosity for *Rec8*, encoding the meiosis-specific kleisin subunit of cohesin, reduced the level of cohesin associated with metaphase-I chromosomes in J-oocytes by 16.6% (fig S2, A and B) and significantly reduced both anaphase-I lagging and aneuploidy (Fig. 2, A–C). Anaphase-I lagging was also significantly reduced when REC8 was partially degraded using either the TRIM-Away (Fig. 2, D and E; and fig S2)(*20*)or dTAG systems (Fig. 2, F and G; fig S2)(*21, 22*). Finally, lagging was also suppressed when proteolytic cleavage of REC8-cohesin was enhanced by increasing separase activity by microinjecting cRNA encoding the AA-separase mutant that lacks inhibitory CDK1 sites (*23*)(Fig. 2, H and I; fig S2). Collectively, these four approaches to weaken cohesion indicate that excessive cohesion at MI impedes homolog segregation in J-oocytes.

**Figure 2.**
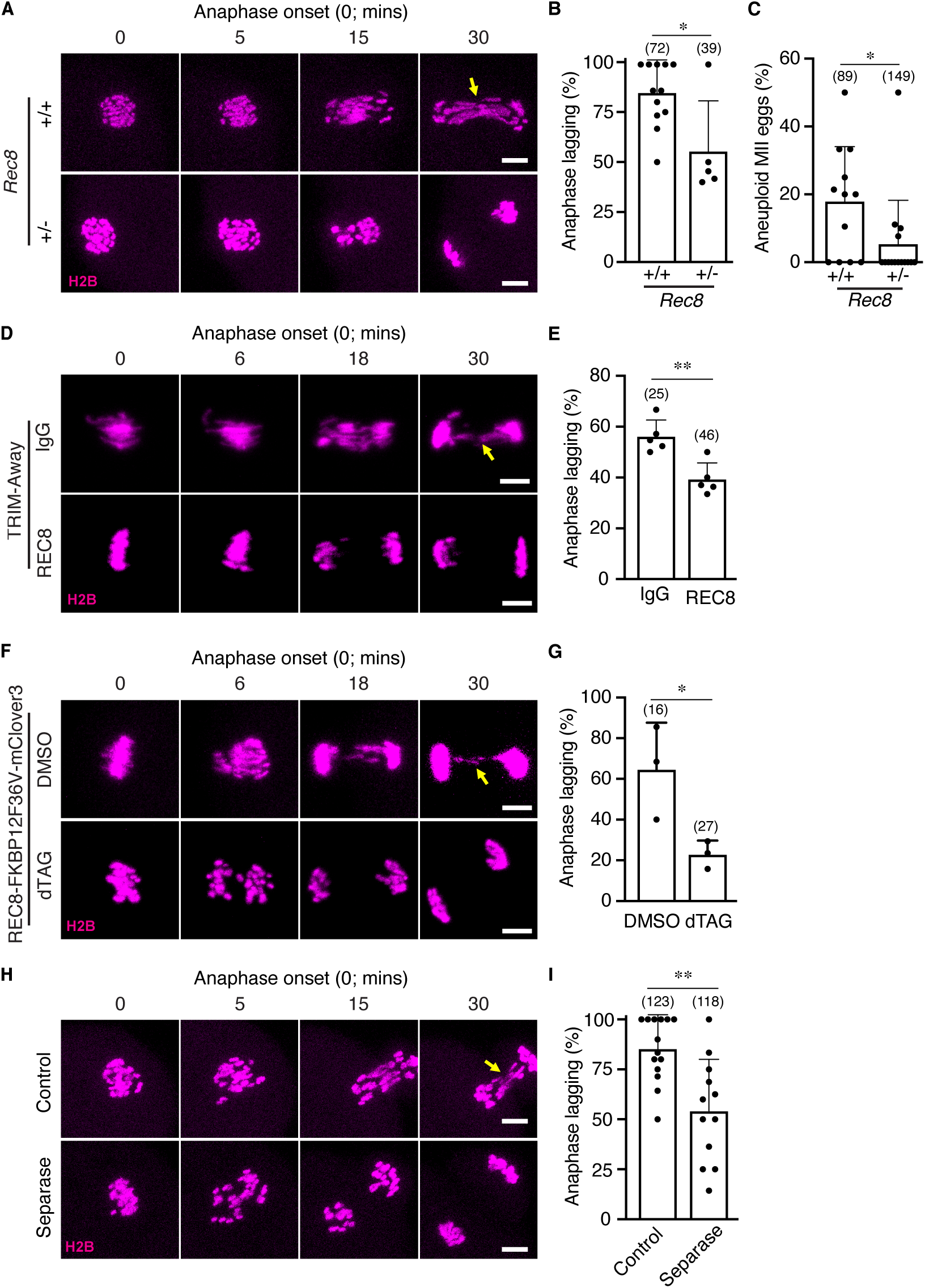
Segregation defects of J-oocytes are rescued by weakening cohesion, or enhancing its resolution during anaphase I. **(A)** Representative frames from live-cell imaging of J-oocytes from *Rec8*^*+/+*^ wild-type and *Rec8*^*+/-*^ heterozygous females. scale bars, 10 µm **(B)** Quantification of chromosome lagging in anaphase I from the experiments represented in A. Each data point represents an individual experiment. Error bars indicate SD. *, *p* <0.05, Mann-Whitney test **(C)** Frequency of aneuploidy detected in metaphase-II chromosome spreads from *Rec8*^*+/+*^ wild-type and *Rec8*^*+/-*^ heterozygous females. *, *p* <0.05, Mann-Whitney test **(D)** Representative frames from live-cell imaging of J-oocytes with (REC8) and without (IgG) partial degradation of REC8 using TRIM-Away (*20*). Scale bars, 10 µm. **(E)** Quantification of chromosome lagging in anaphase I from the experiments represented in D. Each data point represents an individual experiment. Error bars indicate SD. **, *p* <0.01, Unpaired t test **(F)** Representative frames from live-cell imaging of J-oocytes with (dTAG) and without (DMSO) partial degradation of REC8−FKBP12^F36V^−mClover3 using the PROTAC, dTAG-13. (*21, 22*). Scale bars, 10 µm. **(G)** Quantification of chromosome lagging in anaphase I from the experiments represented in F. Each data point represents an individual experiment. Error bars indicate SD. *, *p* <0.05, Unpaired t test **(H)** Representative frames from live-cell imaging of J-oocytes with (separase) and without (control) enhanced separase activity via microinjection of cRNA encoding the AA-separase mutant (*23*). Scale bars, 10 µm. **(I)** Quantification of chromosome lagging in anaphase I from the experiments represented in H. Each data point represents an individual experiment. Error bars indicate SD. **, *p* <0.01, Unpaired t test The yellow arrows in A, D, F, and H highlight lagging chromosomes.

Chromosome lagging is rare and segregation is error-free in Y-oocytes (Fig. 1, A-D) suggesting that cohesion is weaker or more efficiently cleaved than in J-oocytes. Consistent with the former possibility, live-cell imaging of REC8-mClover3 (Fig. 3, A and B) and immunofluorescence staining of REC8 (Fig. 3, C and D; fig. S3A and B, and S4) or SMC3 (a second cohesin subunit, fig S3C and D) revealed that less cohesin was bound to the metaphase-I chromosomes of Y-oocytes relative to those of J-oocytes. Yakoubi et al. recently showed that homolog segregation in a *Mus musculus* x *Mus spicilegus*is hybrid mice is impeded due to mislocalization of the cohesin protector SGO2 to chromosome arms and overprotection of cohesin from separase {El Yakoubi, 2026 #3783}. To explore the possibility that SGO2 is mislocalized in J-oocytes and contributes to lagging and nondisjunction, metaphase-I chromosome spreads were immunostained for SGO2 fig. S3E). However, SGO2 remained restricted to centromeres in J-oocytes.

**Figure 3.**
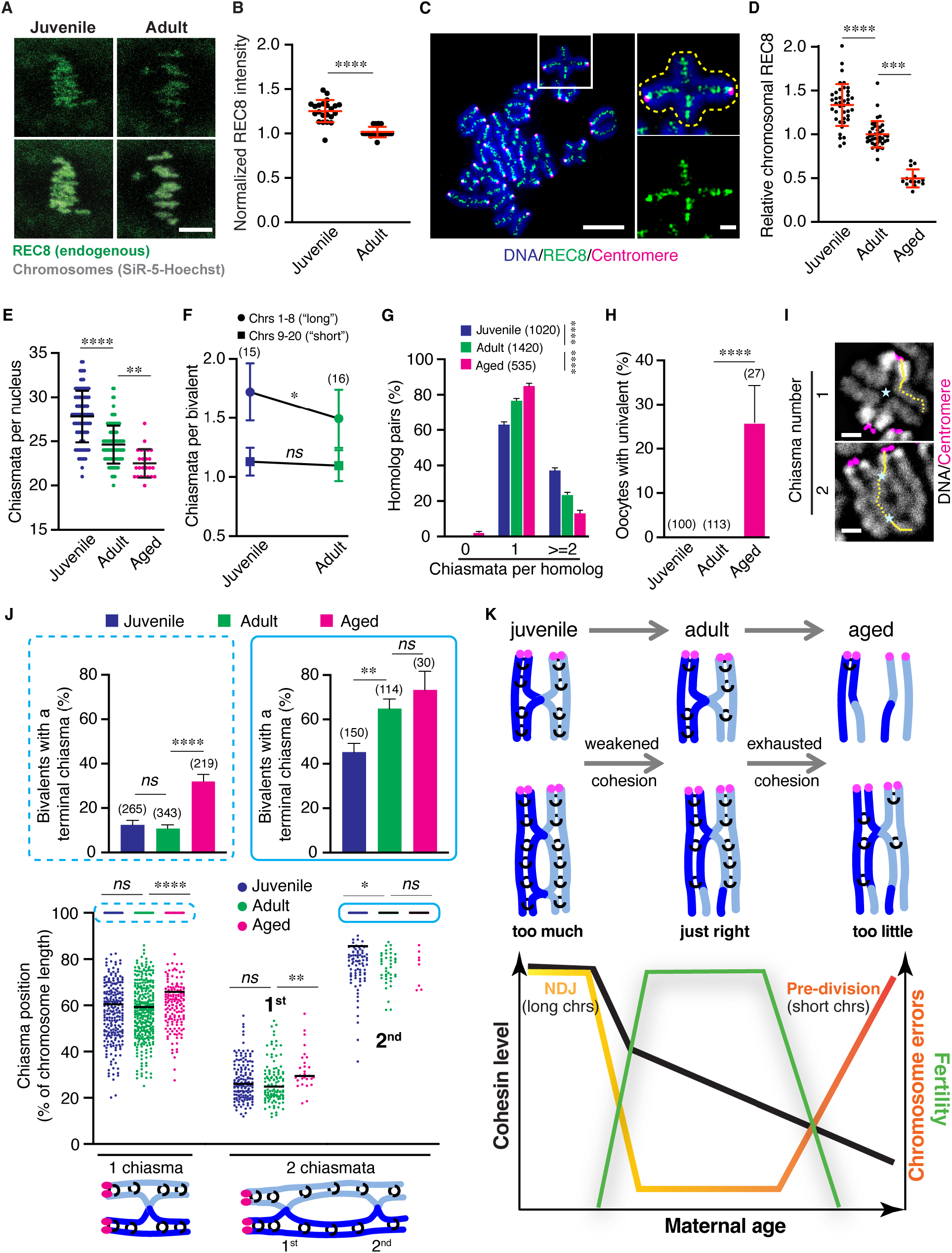
Cohesion strength underlies the frequency and type of segregation errors seen in eggs across the female reproductive lifespan. **(A)** Representative frames from live-cell imaging of J- and Y-oocytes comparing endogenous REC8 levels. The top row shows the REC8-mClover3 channel alone; and the second row shows a merge of REC8-mClover3 and DNA (stained with SiR-DNA). Scale bar, 5 µm **(B)** Quantification of REC8 intensity from the experiments represented in A. Error bars indicate SD. ****, *p* <0.0001; Unpaired t test **(C)** Representative images of a metaphase I oocyte chromosome spread immunstained for REC8 and CREST (centromere) and stained with DAPI (DNA). A single bivalent is magnified showing the merge and REC8 channel alone; and highlighting the ROI used to quantify REC8 signal. scale bar, 10 µm; inset = 2 µm **(D)** Quantification of REC8 intensity from the experiments represented in C. in J-, Y-, and A-oocytes. ***, *p* <0.001; ****, *p* <0.0001; Kruskal-Wallis test with Dunn’s multiple comparisons **(E)** Quantification of chiasmata per nucleus from metaphase-I chromosome spreads from J-, Y-, and A-oocytes. **(F)** Numbers of chiasmata per bivalent for the longest 8 chromosomes (long) versus the shorter 12 chromosomes in individual metaphase-I oocyte chromosomes spreads. Means ± SD are shown. *, *p* <0.05, Unpaired t test **(G)** Percentages of homologs with 0, 1 or ≥2 chiasmata in oocyte metaphase-I chromosome spreads from J-, Y-, and A-oocytes. ****, *p* <0.0001; Fisher’s exact test **(H)** Percentages of oocyte metaphase-I chromosome spreads with unconnected univalent chromosomes from J-, Y-, and A-oocytes. ****, *p* <0.0001; Fisher’s exact test **(I)** Images of individual metaphase-I bivalents immunostained with CREST (centromeres) and stained with DAPI (DNA) connected by one (top) or two (bottom) chiasmata (blue asterisks) and illustrating measurements of chiasma positions (yellow dashed lines). Scale bar, 2 µm **(J)** Lower graphs show positions of chiasmata relative to the centromere of individual bivalents in oocyte metaphase-I chromosome spreads from J-, Y-, and A-oocytes. Positions for bivalents with one or two chiasmata are plotted separately. Data points at 100% of the chromosome length represent very terminal chiasmata. The frequencies of these events are plotted in the upper graphs. For bivalents with a single chiasma, terminalization is only seen in A-oocytes from older animals. For bivalents with two chiasmata, terminalization of the distal chiasmata is already apparent in Y-oocytes from young adults. *, *p* <0.05; **, *p* <0.01; ****, *p* <0.0001; Kruskal-Wallis test with Dunn’s multiple comparisons (for lower graphs) or Fisher’s exact test (for upper graphs) **(K)** Summary of cohesin dynamics and cohesion strength for short and long chromosomes in J-, Y-, and A-oocytes. The graph illustrates frequencies and type of chromosomal errors (yellow-orange line) across the reproductive lifespan as a function of cohesin level (black line). Relative fertility is shown in green. Too much cohesion in J-oocytes impedes homolog segregation, especially for longer chromosomes, leading to nondisjunction. Relatively abrupt weakening of cohesion during sexual maturation remedies these defects and homologs segregate efficiently in young adults. Progressive weakening of cohesion with advancing maternal age eventually hits a critical threshold that results in pre-division errors in which sister-chromatids prematurely separate (*12, 21*). Shorter chromosomes are impacted earlier by this cohesion exhaustion.

Evidence that cohesion is tangibly weaker in Y-oocytes was obtained by demonstrating that centromere-distal chiasmata undergo terminalization (Fig. 3, E-J). Metaphase-I J-oocytes had 27.8 ± 2.9 chiasmata per nucleus, closely matching the number of crossovers formed in prophase-I, based on counts of crossover-specific MLH1 foci (fig. S5; 27.8 ± 2.9 chiasmata vs. 28.6 ± 3.1 MLH1 foci per nucleus, *p*=0.7595). In contrast, Y-oocytes had 24.6 ± 2.1 chiasmata per nucleus (Fig. 3E and fig. S5, A-C; *p*<0.0001 compared to chiasmata numbers in J-oocytes; and *p*<0.0001 compared to MLH1 foci in prophase-I oocytes). Notably, chiasmata were lost specifically from longer chromosomes (Fig. 3F) such that homologs connected by a single chiasma were increased at the expense those with two or more chiasmata (Fig. 3G). However, unconnected univalents were never observed in Y-oocytes (Fig. 3H). Cohesion weakening in J-oocytes was not peculiar to the C57BL/6 mouse background used here for most experiments, as similar results were obtained for oocytes from the PWD and CAST backgrounds (Figs S3, S4, S5). By comparison, A-oocytes from reproductively advanced females showed more severe reductions of both cohesin and chiasmata that were associated with the appearance of univalents in 26.0% of cells (Fig. 3D-H and fig. S3, S4, S5).

These observations are consistent with the idea that weakened cohesion in Y-oocytes manifests as terminalization of the most centromere-distal chiasmata in longer bivalents with two or more chiasmata. The stability of distal chiasmata depends on the small amount of cohesion between the exchange point and the telomere. As such, they are predicted to be the most vulnerable to loss when cohesion is weakened. Consistently, in bivalents with two chiasmata, the “second” centromere-distal chiasma was shifted to a more terminal position in Y-oocytes relative to J-oocytes (Fig. 3J, right-hand side; *p*=0.0228, Kruskal-Wallis test with Dunn’s multiple comparisons). The average positions of centromere-proximal chiasmata did not change. Similarly, in bivalents with a single chiasma, average chiasma position was not different between J- and Y-oocytes (Fig. 3J, left-hand side). By contrast, the greater loss of cohesion in A-oocytes was associated with more terminal locations of all chiasmata.

Our findings suggest that high levels of aneuploidy in the eggs of adolescent females is a conserved feature of mammalian oogenesis (*18, 19*). We define the underlying cause as being excessive cohesion that is inefficiently resolved at anaphase-I, such that homolog segregation is impeded. Consistently, long chromosomes, which have more cohesin, are more prone to nondisjoin in the eggs of adolescent women {Gruhn, 2019 #3553}. In adults, this defect is remedied and homolog disjunction is efficient because cohesin levels are lower and cohesion is demonstrably weaker. The surprising finding that cohesion is already being weakened during sexual maturation, and not just during reproductive aging, defines a Goldilocks scenario for egg quality in which cohesion strength underlies the frequency and type of segregation errors across the female reproductive lifespan (Fig. 3K). Future studies aim to understand what limits the efficiency of cohesin cleavage in anaphase-I J-oocytes, the mechanism of fractional cohesin loss in Y-oocytes, and how this loss is balanced so that that segregation is efficient, but bivalents remain intact. Our findings have implications for fertility treatments and *in vitro* fertilization using eggs from adolescent women, suggesting that weakening cohesion could increase the chances of obtaining viable euploid eggs.

## Acknowledgments

We thank Michael Lampson (University of Pennsylvania) and Scott Keeney (Memorial Sloan-Kettering Cancer Center) for reagents, and members of the Hunter, Yun and Mogessie labs for discussions.

## Funding

Eunice Kennedy Shriver National Institute of Child Health and Human Development grant 5R01HD109322 (NH)

Guangdong Basic and Applied Basic Research Foundation 2024A1515012907 and 2023A1515220243 (YY)

Vallee Scholars Award (BM), VS-2024-56 Pew Scholars Award (BM), 00037689

National Institutes of Health (BM), R35GM146725 Lalor Foundation Postdoctoral Fellowship (JL)

NH is an Investigator with the Howard Hughes Medical Institute

## Author contributions

Conceptualization: YY, NH

Methodology: YY, NH, BM, JL, HB

Investigation: YY, KI, DD, ZG, JL, HB

Funding acquisition: NH, YY, BM

Project administration: YY, NH

Supervision: NH, BM

Writing – original draft: YY, NH

Writing – review & editing: YY, NH, BM, KI, DD, ZG, JL, HB

## Data and materials availability

All other data are available in the main text or the supplementary materials.

## Supplementary Materials

Materials and Methods Figs. S1 to S5

References

Movies S1 and S2

## References and Notes

1. E. E. Telfer, J. Grosbois, Y. L. Odey, R. Rosario, R. A. Anderson, Making a good egg: human oocyte health, aging, and in vitro development. Physiol Rev 103, 2623–2677 (2023).

2. M. Hu et al., PRC1-mediated epigenetic programming is required to generate the ovarian reserve. Nature communications 13, 4510 (2022).

3. N. Hunter, Oocyte Quality Control: Causes, Mechanisms, and Consequences. Cold Spring Harb Symp Quant Biol 82, 235–247 (2017).

4. N. Hunter, Meiotic Recombination: The Essence of Heredity. Cold Spring Harbor perspectives in biology 7, (2015).

5. K. I. Ishiguro, The cohesin complex in mammalian meiosis. Genes to cells : devoted to molecular & cellular mechanisms 24, 6–30 (2019).

6. V. Makrantoni, A. L. Marston, Cohesin and chromosome segregation. Current biology : CB 28, R688–R693 (2018).

7. Y. Murayama, C. P. Samora, Y. Kurokawa, H. Iwasaki, F. Uhlmann, Establishment of DNA-DNA Interactions by the Cohesin Ring. Cell 172, 465–477 e415 (2018).

8. K. Wassmann, Separase Control and Cohesin Cleavage in Oocytes: Should I Stay or Should I Go? Cells 11, (2022).

9. R. Jessberger, Age-related aneuploidy through cohesion exhaustion. EMBO reports 13, 539–546 (2012).

10. W. Huang, X. Li, H. Yang, H. Huang, The impact of maternal age on aneuploidy in oocytes: Reproductive consequences, molecular mechanisms, and future directions. Ageing Res Rev 97, 102292 (2024).

11. L. Wartosch et al., Origins and mechanisms leading to aneuploidy in human eggs. Prenat Diagn 41, 620–630 (2021).

12. T. Chiang, F. E. Duncan, K. Schindler, R. M. Schultz, M. A. Lampson, Evidence that weakened centromere cohesion is a leading cause of age-related aneuploidy in oocytes. Current biology : CB 20, 1522–1528 (2010).

13. L. M. Lister et al., Age-related meiotic segregation errors in mammalian oocytes are preceded by depletion of cohesin and Sgo2. Current biology : CB 20, 1511–1521 (2010).

14. Y. Yun, S. I. Lane, K. T. Jones, Premature dyad separation in meiosis II is the major segregation error with maternal age in mouse oocytes. Development 141, 199–208 (2014).

15. B. P. Mihalas et al., Age-dependent loss of cohesion protection in human oocytes. Current biology : CB 34, 117–131 e115 (2024).

16. F. E. Duncan et al., Chromosome cohesion decreases in human eggs with advanced maternal age. Aging Cell 11, 1121–1124 (2012).

17. J. A. Merriman, P. C. Jennings, E. A. McLaughlin, K. T. Jones, Effect of aging on superovulation efficiency, aneuploidy rates, and sister chromatid cohesion in mice aged up to 15 months. Biology of reproduction 86, 49 (2012).

18. J. R. Gruhn et al., Chromosome errors in human eggs shape natural fertility over reproductive life span. Science 365, 1466–1469 (2019).

19. D. Lechniak et al., Gilts and sows produce similar rate of diploid oocytes in vitro whereas the incidence of aneuploidy differs significantly. Theriogenology 68, 755–762 (2007).

20. D. Clift, C. So, W. A. McEwan, L. C. James, M. Schuh, Acute and rapid degradation of endogenous proteins by Trim-Away. Nat Protoc 13, 2149–2175 (2018).

21. J. Leem et al., A versatile cohesion manipulation system probes female reproductive age-related egg aneuploidy. Nat Aging 5, 2215–2227 (2025).

22. K. Li, C. M. Crews, PROTACs: past, present and future. Chem Soc Rev 51, 5214–5236 (2022).

23. T. Chiang, R. M. Schultz, M. A. Lampson, Age-dependent susceptibility of chromosome cohesion to premature separase activation in mouse oocytes. Biology of reproduction 85, 1279–1283 (2011).

