## Supplementary Materials for "A cohesion optimum underlies chromosome segregation fidelity in oocytes"

Materials and Methods
Figs. S1 to S5
Supplementary references
Movies S1 and S2

### Materials and Methods

#### Animals

Wild-type mouse strains C57BL/6J (Stock No. 000664), PWD/PhJ (Stock No. 004660), and CAST/EiJ (Stock No. 000735) were obtained from The Jackson Laboratory. The B6.Cg-Rec8$^{mei8}$/JcsMmjax strain (MMRRC_034762-JAX) was acquired from the Mutant Mouse Resource and Research Center (MMRRC) at The Jackson Laboratory, courtesy of Dr. John Schimenti (Cornell University) (*1*). The REC8−FKBP12$^{F36V}$−mClover3 mouse model was generated via CRISPR−Cas9-mediated genome editing as previously described (*2*).

Animals were housed in groups of up to four per cage in a temperature-controlled vivarium under a 12-h light/dark cycle, with ad libitum access to standard rodent chow and water. All animal procedures were approved by the Institutional Animal Care and Use Committees (IACUC) of the University of California, Davis (approval number 2026-24835), and Yale University (2021-20408), and were performed in accordance with their guidelines.

#### Surface Spreading and Immunofluorescence Staining of Prophase I Fetal Oocyte Nuclei

Ovaries from E18.5 female fetuses were dissected. Surface spreading of prophase I oocyte nuclei was performed as described (*3*). Briefly, ovaries were incubated on ice for 20 min in hypotonic extraction buffer, then minced in 20 µL 0.1 M sucrose (Sigma-Aldrich, S0389). Cell suspension was spread onto slides (Fisher Scientific, 12-544-7) pre-coated with 1% PFA / 0.15% Triton X-100 (pH 9.2) and air-dried overnight in a humid chamber. Slides were washed in 0.4% Photo Flo (Kodak, 1464510) and air-dried.

For immunofluorescence, slides were blocked 1 h (RT), then incubated overnight (RT) with primary antibodies: rabbit anti-SYCP3 (Santa Cruz, sc-33195; 1:200), mouse anti-MLH1 (Cell Signaling Technology, 3515; 1:25), human anti-centromere (ACA/CREST; ImmunoVision, HCT-0100; 1:1000). After washing, slides were incubated 1 h (RT) with secondary antibodies: goat anti-rabbit 568 (Thermo Fisher Scientific; A11036; 1:1000), goat anti-mouse 488 (Thermo Fisher Scientific; A11029; 1:1000), and goat anti-human DyLight 649 (Jackson Labs, 109-495-088; 1:200). Coverslips were mounted with ProLong Diamond (Thermo Fisher Scientific, P36970) prior to imaging. Images were acquired using a Zeiss AxioPlan II microscope equipped with a 63X oil immersion objective and Hamamatsu ORCA-ER CCD camera.

#### Isolation and In Vitro Maturation of Fully Grown GV-Stage Oocytes

Germinal vesicle (GV)-stage oocytes were collected from mice that had been hormonally primed with an intraperitoneal injection of 5 IU pregnant mare serum gonadotropin (PMSG; Sigma-Aldrich, G4877). Oocyte collection was performed by multiple punctures of ovarian follicles with a 25G needle under a stereo-microscope (Nikon, SMZ1500). Only oocytes surrounded by an intact cumulus cell layer were selected. All manipulations were carried out in M2 medium (Sigma-Aldrich, M7167) supplemented with 2.5 µM milrinone at 37 °C to maintain meiotic arrest. Meiotic maturation was stimulated by milrinone washout. Prior to maturation culture, cumulus cells were mechanically removed to allow subsequent observation of germinal vesicle breakdown (GVBD) and polar body extrusion (PBE). Only oocytes that underwent GVBD within 3 h were included for quantification of PBE after an additional 13 h of culture. All cultures were performed under mineral oil (Nidacon, NO-100).

**Chromosome Spreading of Metaphase I Oocytes and Immunofluorescence Staining**

Oocytes exhibiting GVBD within 60 ± 15 min of culture were selected, and metaphase I oocytes were subsequently obtained after a total of 7 h of culture. Metaphase-I chromosome spreads were prepared as described (*3*). Briefly, the zona pellucida was removed with Acid Tyrode's solution (Sigma-Aldrich, M1788). Zona-free oocytes were spread onto glass slides in 1% paraformaldehyde (Electron Microscopy Science; 19208) containing 0.15% Triton X-100 (Sigma-Aldrich, X100) and 3 mM DTT (USBiological, D8070) (pH 9.2). To facilitate accurate chromosome counting, each oocyte was processed in an individual well of a 12-well slide (Electron Microscopy Science; 63425-05). Chromosome spreads were air-dried overnight at RT.

All subsequent steps were performed at RT. Slides were blocked for 1–2 h in blocking solution, then incubated overnight with primary antibodies: human anti-centromere (ACA/CREST; ImmunoVision; HCT-0100; 1:1000), mouse anti-TRF1 (Abcam; ab10579; 1:100), together with either rabbit anti-REC8 (gift from Dr. Scott Keeney; 1:100), rabbit anti-SMC3 mAb (ABclonal; A19591; 1:100), or rabbit anti-SGO2 pAb (Abmart; PC24137; 1:100). After three washes in blocking solution, slides were incubated for 1 h with secondary antibodies (Thermo Fisher Scientific): goat anti-human 555 (A-21433; 1:1000) and goat anti-rabbit 488 (A-11034; 1:1000). Chromosomes were counterstained with DAPI (5 µg/mL; Sigma-Aldrich; D8417). Images were acquired on a Zeiss AxioPlan II microscope with a 63× oil objective and a Hamamatsu ORCA-ER CCD camera.

**Chromosome Spreading of Metaphase II Eggs and Immunofluorescence Staining**

Metaphase II eggs were collected after 16 h of culture, and chromosome spreading was performed as described for metaphase I oocytes. To facilitate subsequent analysis of aneuploidy, each egg and its corresponding first polar body were spread into separate wells of a 12-well slide (Electron Microscopy Science; 63425-05).

**cRNA Synthesis and Microinjection of GV-Stage Oocytes**

The following plasmids were used for cRNA synthesis, as previously described: pCS2-H2B-mCherry (Addgene; #135604) (*4*), pGEMHE-H2B-mEGFP (Addgene; #105528) (*5*), pIVT-AA-separase (gift from Dr. Michael Lampson) (*6*), and pGEM-SNAP-TRIM21 (*7*). 5′-capped cRNAs were synthesized using the mMESSAGE mMACHINE™ T7 ULTRA Transcription Kit (Thermo Fisher Scientific/Ambion; AM1345) and dissolved in nuclease-free water. mRNA concentrations were measured with a NanoDrop spectrophotometer (Thermo Fisher Scientific).

cRNA microinjection was performed as described previously (*8*) using an Eppendorf microinjection system (FEMTOJET 4I) mounted on a heated stage of a Nikon Ti2-A inverted microscope. Approximately 1–3 pL of cRNA solution was injected into the cytoplasm of GV-stage oocytes at the following pipette concentrations: 50 ng/µL for H2B-mCherry, 50 ng/µL for H2B-mEGFP, 50 ng/µL for TRIM21, 100 ng/µL for AA-separase (rescue experiment), and 2000 ng/µL for AA-separase (overexpression purpose).

**TRIM-Away-Mediated REC8 Partial Degradation in Juvenile Mouse Oocytes**

For partial REC8 degradation using TRIM-Away, fully grown GV-stage oocytes isolated from 3-week-old juvenile mice were co-microinjected with a mixture containing H2B-mCherry cRNA,

TRIM21 cRNA, and REC8 antiserum (a gift from Dr. Michael Lampson; 1:1000 dilution) or control IgG, at a total volume of 2–3 pL. Chromosomes were fluorescently labeled by H2B-mCherry to enable subsequent live-cell imaging.

**Confocal Live-Cell Imaging**

With the exception of the endogenous REC8 signal live-imaging experiment (Fig. 3A), all chromosomes were fluorescently labeled with H2B-mCherry or H2B-mEGFP. High-resolution live-cell imaging was performed using a Zeiss LSM 800 confocal microscope equipped with an Airyscan detector, a 40× water-immersion objective, an Axiocam camera, and an environmental chamber maintained at 37 °C. Chromosomes were imaged with 18 z-sections at 2.0-μm interval. For chromosome dynamics analysis, sequential image stacks were acquired every 5 or 6 min. To capture chromosome alignment and segregation events, imaging was initiated upon milrinone washout and continued for up to 24 h of culture.

To induce rapid endogenous REC8 degradation in juvenile oocytes from REC8−FKBP12$^{F36V}$−mClover3 mice, GV-stage oocytes expressing H2B-mCherry were induced to resume meiosis. After 6 h of culture, oocytes were treated with 1 μM dTAG-13 (Merck; SML2601) or DMSO (Merck; D2650-5X5ML) as a control, and immediately subjected to live-cell imaging using the same acquisition parameters.

Metaphase I chromosomes in oocytes from the REC8-FKBP12$^{F36V}$-mClover3 mice were labeled with 100 nM SiR-DNA dye (Cytoskeleton Inc; CY-SC007) diluted in M2 medium. REC8-mClover3 was imaged on a Leica STELLARIS 5 confocal laser scanning microscope equipped with an environmental incubator box and a ×40 C-Apochromat 1.2 numerical aperture water immersion objective. Images were acquired at a temporal resolution of 3 minutes, with Z stacks totaling approximately 40 μm and confocal sections spaced 1.5 μm apart. Oocytes were imaged in M2 medium under mineral oil, as described previously (*9*).

**Quantification of chromosome-associated proteins on metaphase-I spreads**

Image processing and quantification were performed using ImageJ/Fiji (NIH). Metaphase-I spreads from different age groups were prepared and processed in parallel; all images were acquired on the same day with identical exposure settings. For REC8 and SMC3, individual chromosomes were manually selected using the DAPI channel. Mean gray values of REC8/SMC3 and CREST were measured after background subtraction. The relative chromosomal intensity was calculated as the REC8/SMC3-to-CREST ratio, then averaged across all chromosomes per oocyte. For centromere-enriched proteins SGO2, mean fluorescence intensities of SGO2 and CREST were calculated within a 2-μm-diameter circle centered on the CREST signals of each sister-kinetochore pair after background subtraction. The relative level was calculated as the SGO2-to-CREST ratio, then averaged across all sister-kinetochore pairs per oocyte. At least three independent experiments were pooled, with each value normalized to the mean of the corresponding control oocytes.

**Measuring chiasma terminalization on metaphase-I spreads**

All chiasma measurements were performed using ImageJ/Fiji (NIH). For chiasma counting, only spreads in which all 20 bivalents were present and the chromosomes were well spread and non-overlapping were selected for comparison. The length of a chromosome arm was defined as the linear pixel distance from the center of the centromeric CREST signal to the distal termination of the DAPI-stained arm (telomere). For the chromosome length-based grouping shown in Fig. 3F, the arm length of each bivalent within an individual oocyte was used to rank all 20 bivalents. They were then divided into two categories: the eight longest chromosomes ("long") and the remaining twelve chromosomes ("short") per oocyte.

To further quantify chiasma distribution, the position of a chiasma was determined by measuring the distance from the centromere to the centroid of the chiasma constriction. When two chiasmata were present, each was measured individually. The relative chiasma position was calculated as: distance from centromere to chiasma / total arm length. Thus, a value approaching 1 indicates a highly terminalized chiasma near the telomere, whereas a value close to 0 reflects a proximal position. Global terminalization was assessed by comparing the distributions of relative chiasma position values across bivalents from different age groups. In the chiasma distribution assay, any bivalents with ambiguous chiasma configurations were excluded.

**Quantification of Live-Cell Imaging Data**

In live-cell imaging, anaphase onset was defined as the first frame in which the aligned homologous chromosomes at the metaphase plate initiated separation and began moving toward opposite poles. Lagging chromosomes were defined as those that persisted at the metaphase plate for at least 30 min after anaphase onset.

Endogenous REC8 levels were quantified in live oocytes from REC8-FKBP12$^{F36V}$-mClover3 mice after 6 hours of release from GV arrest. High-resolution live-cell imaging datasets were analyzed with Imaris software (Bitplane). To ensure consistent quantification under equivalent imaging conditions and to measure fluorescence at maximal signal intensity, analysis was performed on the first time point of each live-imaging dataset. Chromosomes were segmented using the SiR-DNA fluorescence signal to generate a three-dimensional chromosomal mask. The mean mClover3 fluorescence intensity within the masked chromosomal region was measured to quantify chromosome-associated REC8 levels.

**Statistical Analysis**

Due to the intrinsic limitations of statistical analysis of chromosome errors in metaphase-I and metaphase-II chromosome spreads, and the small number of analyzable nuclei obtainable from individual females, oocyte samples from independent experiments were pooled to calculate proportions. Comparisons of proportions were performed using Fisher's exact test, and error bars for such data represent the standard error of the proportion as previous described (*10*, *11*).

Mean values are presented as mean ± standard deviation (SD). For comparisons between two groups, an unpaired *t*-test (parametric) or Mann–Whitney test (nonparametric) was used.
For multiple comparisons, ordinary one-way ANOVA with post hoc Holm-Šídák or Tukey's test (parametric), or the Kruskal–Wallis test followed by Dunn's post hoc test (nonparametric), was applied.

All data were analyzed using GraphPad Prism 10, with a significance threshold of $p < 0.05$. At least three independent experiments were performed for each comparison. The numbers of experiments and oocytes analyzed are indicated in the figures or their legends.

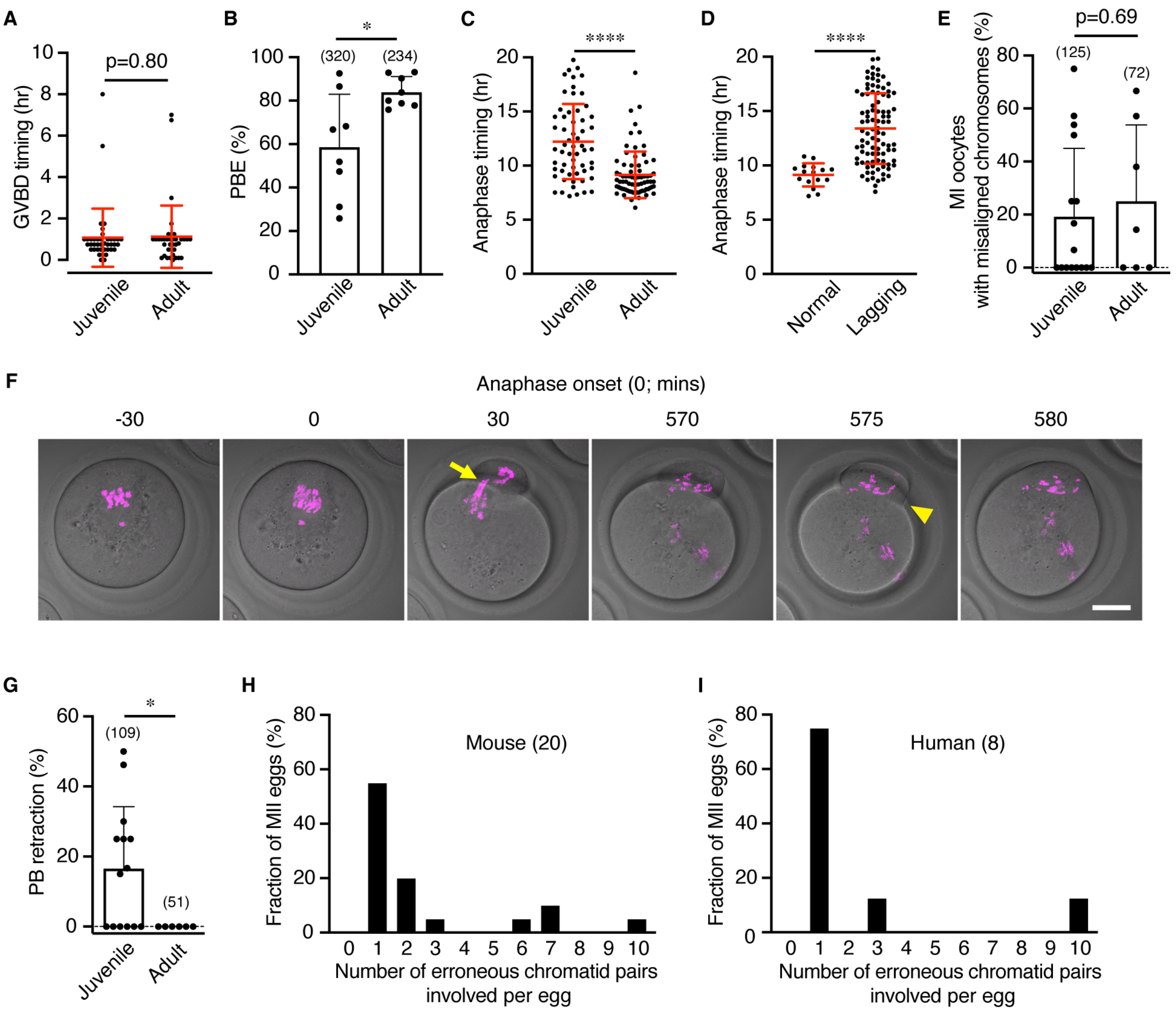


**Supplementary Figure S1 (related to Figure 1)**

(**A**). The timing of germinal vesical breakdown (GVBD, meiosis resumption). Error bars show means ± SD.

(**B**). Efficiency of polar-body extrusion, indicative of successful MI. Error bars show means ± SD. *, $p$<0.05, Unpaired t test

(**C**). The timing of anaphase-I onset (hrs post GVBD). Error bars show means ± SD. ****, $p$<0.0001, Mann-Whitney test

(**D**). Anaphase-I timing in J-oocytes with normal chromosome segregation or lagging. Error bars show means ± SD. ****, $p$<0.0001, Mann-Whitney test

(**E**). Percentage of metaphase-I oocytes with misaligned chromosomes. Error bars show means ± SD.

(**F**). Representative frames from live-cell imaging of  a J-oocyte exhibiting polar body retraction. The arrow highlights lagging chromosomes; the caret indicate the site of polar-body retraction. **What does the caret indicate?** Scale bar, 20 µm

(**G**). Percentage of oocytes exhibiting polar body retraction. Error bars show means ± SD. *, $p$<0.05, Mann-Whitney test

(**H**). Numbers of chromosome errors in individual aneuploid MII eggs from juvenile mice.

(**I**). Numbers of chromosome errors in individual aneuploid MII eggs from adolescent women, (<20 years old/ Adapted from Gruhn et al. (11))

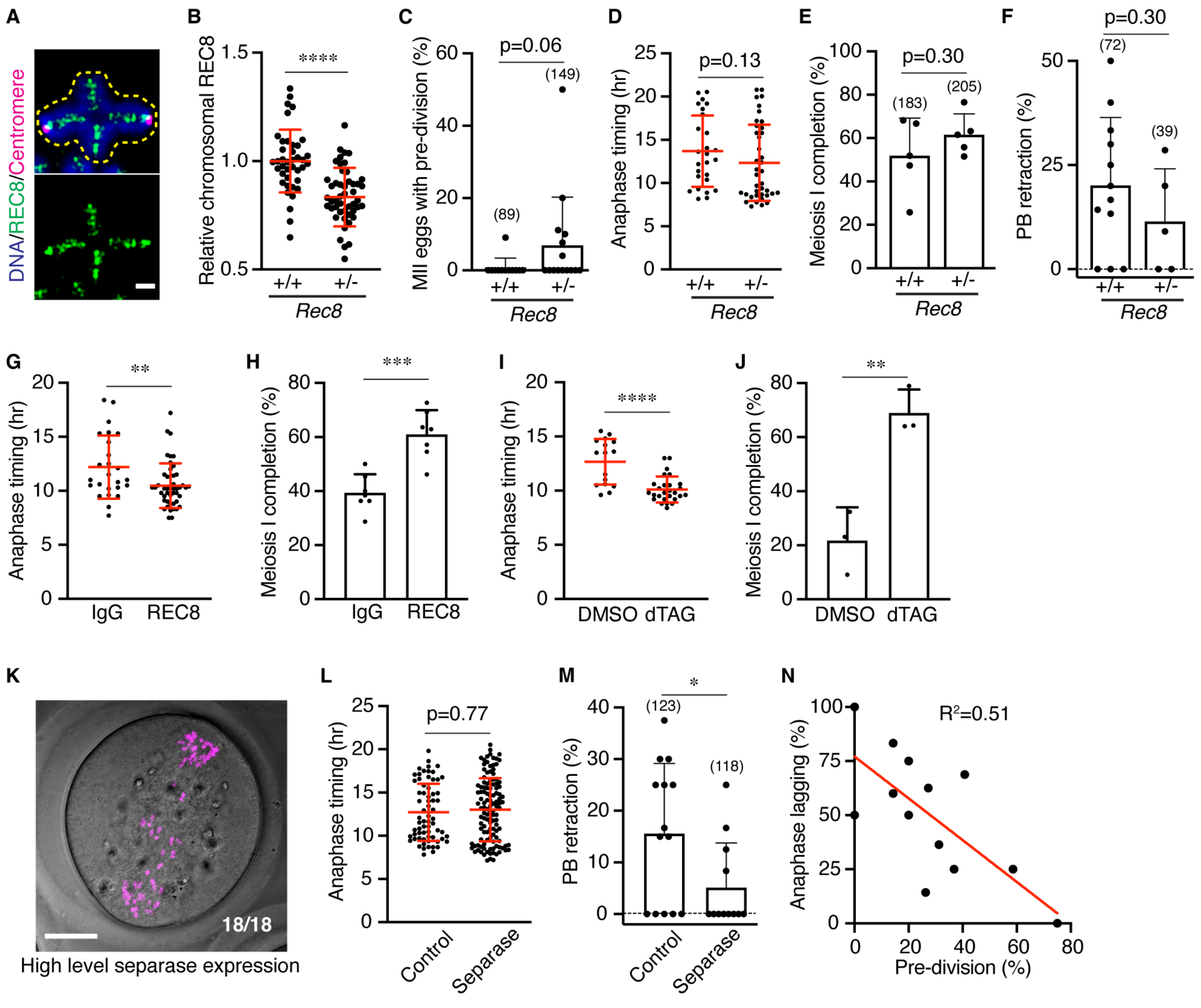


**Supplementary Figure S2 (related to Figure 2)**

(**A**). Representative image of an individual metaphase-I oocyte immunostained for REC8 and CREST (centromere) and stained with DAPI (DNA). The yellow dashed line indicates the ROI used to quantify REC8 signal. Scale bar, 2 μm.

(**B**). Quantification of REC8 intensity on metaphase-I J-oocytes from *Rec8*$^{+/+}$ wild-type and *Rec8*$^{+/-}$ heterozygous females. Error bars show means ± SD. ****, $p$<0.0001, Unpaired t test

(**C**). Frequency of pre-division errors (single chromatids) in metaphase-II J-oocytes suggests slightly elevated separation of sister-chromatids when *Rec8* is heterozygous. Error bars show means ± SD.

(**D**). Anaphase-I timing in J-oocytes from *Rec8*$^{+/+}$ wild-type and *Rec8*$^{+/-}$ heterozygous females. Error bars show means ± SD.

(**E**). Rate of Meiosis I completion in J-oocytes from *Rec8*$^{+/+}$ wild-type and *Rec8*$^{+/-}$ heterozygous females suggest a higher efficiency when *Rec8* is heterozygous. Error bars show means ± SD.

(**F**). Rate of polar-body retraction in J-oocytes from *Rec8*$^{+/+}$ wild-type and *Rec8*$^{+/-}$ heterozygous females suggests a lower rate when *Rec8* is heterozygous. Error bars show means ± SD.

(**G**). Anaphase-I timing in J-oocytes with (REC8) and without (IgG) partial degradation of REC8 using TRIM-Away indicates faster timing when REC8 levels are lower. Error bars show means ± SD. **, $p<0.01$, Mann-Whitney test

(**H**) Rate of Meiosis I completion in J-oocytes with (REC8) and without (IgG) partial degradation of REC8 using TRIM-Away indicates a higher efficiency when REC8 levels are lower. Error bars show means ± SD. ***, $p<0.001$, Unpaired t test

(**I**). Anaphase-I timing in J-oocytes with (dTAG) and without (DMSO) partial degradation of REC8−FKBP12$^{F36V}$−mClover3 using the PROTAC, dTAG-13 indicates faster timing when REC8 levels are lower. Error bars show means ± SD. ****, $p<0.0001$, Mann-Whitney test

(**J**). Rate of Meiosis I completion in J-oocytes with (dTAG) and without (DMSO) partial degradation of REC8−FKBP12$^{F36V}$−mClover3 using the PROTAC, dTAG-13 indicates a higher efficiency when REC8 levels are lower. Error bars show means ± SD. **, $p<0.01$, Unpaired t test

(**K**). Representative image of a J-oocyte with a high level of AA-separase expression (high concentration cRNA injection) showing high levels of premature sister-chromatid separation. Scale bar, 20 µm

(**L**). Anaphase-I timing in J-oocytes with (separase) and without (control) enhanced separase activity via microinjection of AA-separase cRNA shows that anaphase-I is not accelerated. Error bars show means ± SD.

(**M**). Rate of polar-body retraction in J-oocytes with (separase) and without (control) enhanced separase activity indicates a lower rate when separase activity is higher. Error bars show means ± SD. *, $p<0.05$, Mann-Whitney test

(**N**). Correlation between the fractions of oocytes with anaphase lagging and pre-division in J-oocytes injected with AA-separase cRNA. A high level of exogenous separase activity, indicated by a high percentage of oocytes with evidence of sister-chromatid separation (pre-division) correlates with a lower frequency of anaphase lagging.

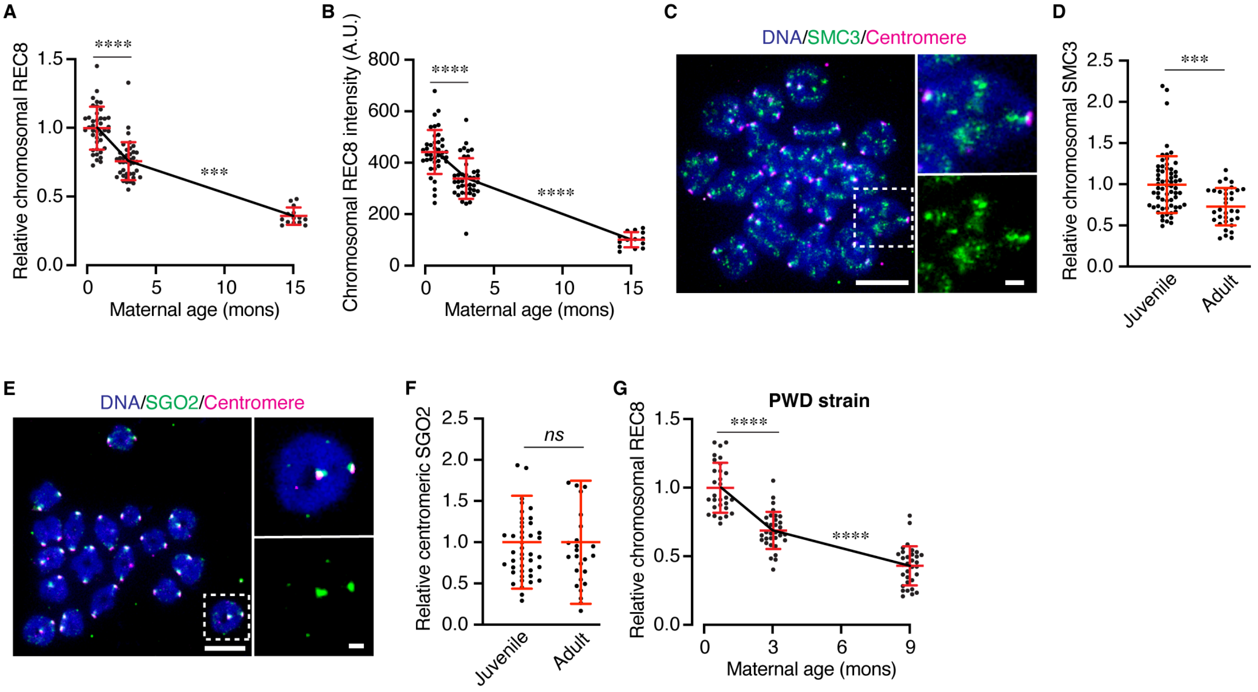


**Supplementary Figure S3 (related to Figure 3)**

(**A**). Chronological plots of relative REC8 intensity (REC8/CREST ratio) measured from metaphase-I oocytes. Error bars show means ± SD. ***, $p$<0.001; ****, $p$<0.0001; Kruskal-Wallis test with Dunn's multiple comparisons

(**B**). Chronological plots absolute REC8 intensity measured from metaphase-I oocytes. Error bars show means ± SD. ****, $p$<0.0001; Ordinary one-way ANOVA with Holm-Šídák's multiple comparisons test

(**C**). Representative image of a metaphase I oocyte chromosome spread immunostained for SMC3 and CREST (centromere) and stained with DAPI (DNA). A single bivalent is magnified showing the merge and SMC3 channel alone. Scale bar, 10 µm; inset = 2 µm

(**D**). Quantification of relative SMC3 intensity from the experiments represented in C. Error bars show means ± SD. ***, $p$<0.001; ****, Unpaired t test

(**E**). Representative image of a metaphase I J-oocyte chromosome spread immunostained for SGO2 and CREST (centromere), and stained with DAPI (DNA). SGO2 does not mislocalize to chromosome arms in J-oocytes. Scale bar, 10 µm; inset = 2 µm

(**F**). Quantification of relative SGO2 intensity show that levels are not different between J- and Y-oocytes. Error bars show means ± SD.

(**G**). Chronological plots of relative REC8 intensity measured from metaphase-I oocytes from the PWD mouse strain. Error bars show means ± SD. ****, $p<0.0001$; Ordinary one-way ANOVA with Tukey's multiple comparisons test

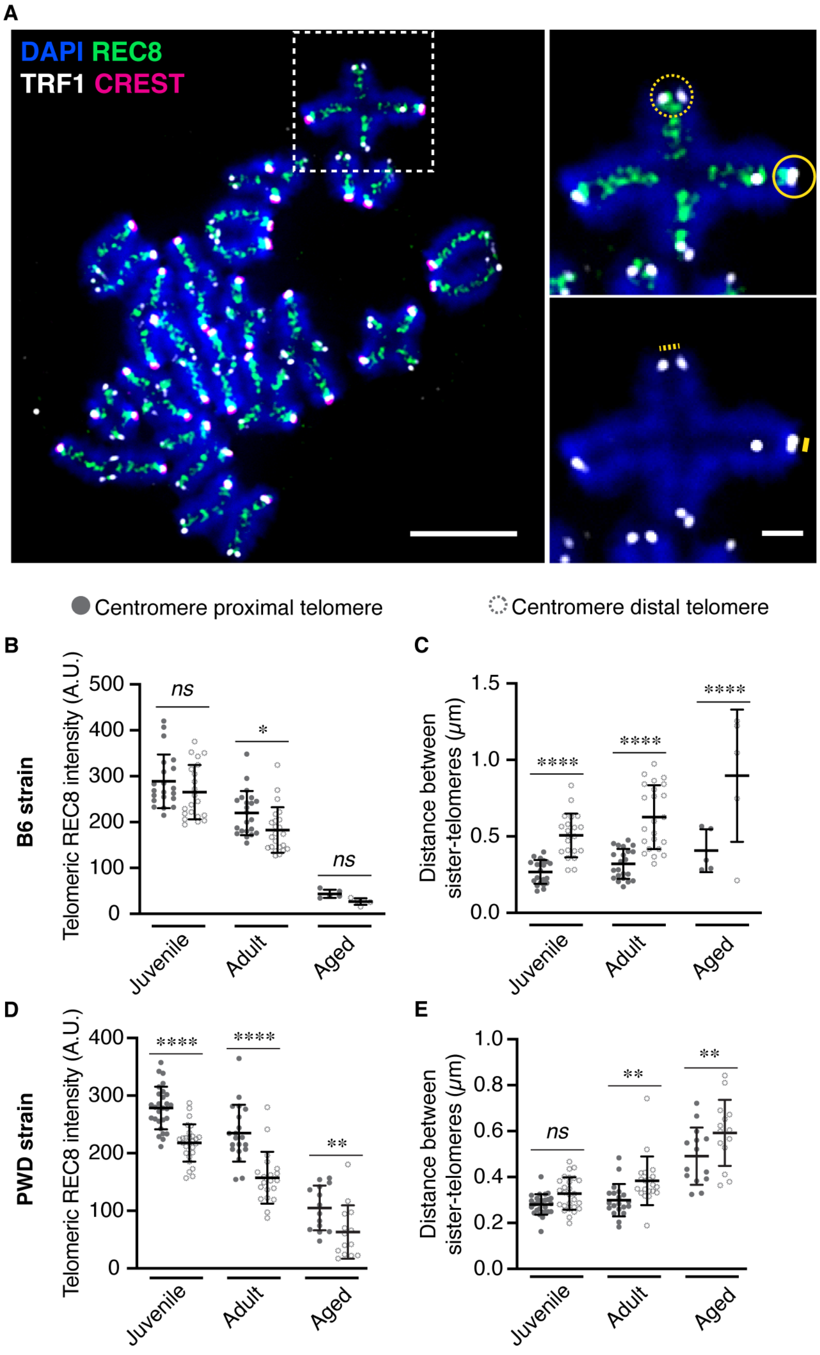


**Supplementary Figure S4 (related to Figure 3)**

(**A**). Representative image of a metaphase-I oocyte chromosome spread immunostained for the telomere marker, TRF1, REC8 and CREST (centromere) and stained with DAPI (DNA). A single bivalent is magnified showing the merge of TRF1+REC8+DNA channels (top) and the TRF1+DAPI channels (bottom). At the centromere distal telomeres (no CREST staining), the dashed circle indicates the ROI used to quantify REC8; and the dashed line indicates the distance between sister telomeres. At the centromere proximal telomeres (with CREST staining), the solid circle indicates the ROI used to quantify REC8; and the solid line indicates the distance between sister telomeres. Scale bar, 10 µm; inset = 2 µm

(**B**). Comparisons of telomeric REC8 intensity from the experiments represented in A show that more REC8 is present at the centromere-proximal versus distal telomeres, but that the levels reduce proportionally with age. Error bars show means ± SD. *, $p<0.05$; Two-way ANOVA with multiple comparisons

(**C**). Comparisons of distances between sister-telomeres from the experiments represented in A show that sister telomeres are further apart the centromere-distal end consistent with weaker cohesion. Error bars show means ± SD. ****, $p<0.0001$; Two-way ANOVA with multiple comparisons

(**D**). Comparisons of telomeric REC8 intensity as in panel B, but for oocytes from the PWD mouse strain. **, $p<0.01$; ****; $p<0.0001$; Two-way ANOVA with multiple comparisons

(**E**). Comparisons of distances between sister-telomeres as in panel C but for oocytes from the PWD mouse strain. **, $p<0.01$; Two-way ANOVA with multiple comparisons

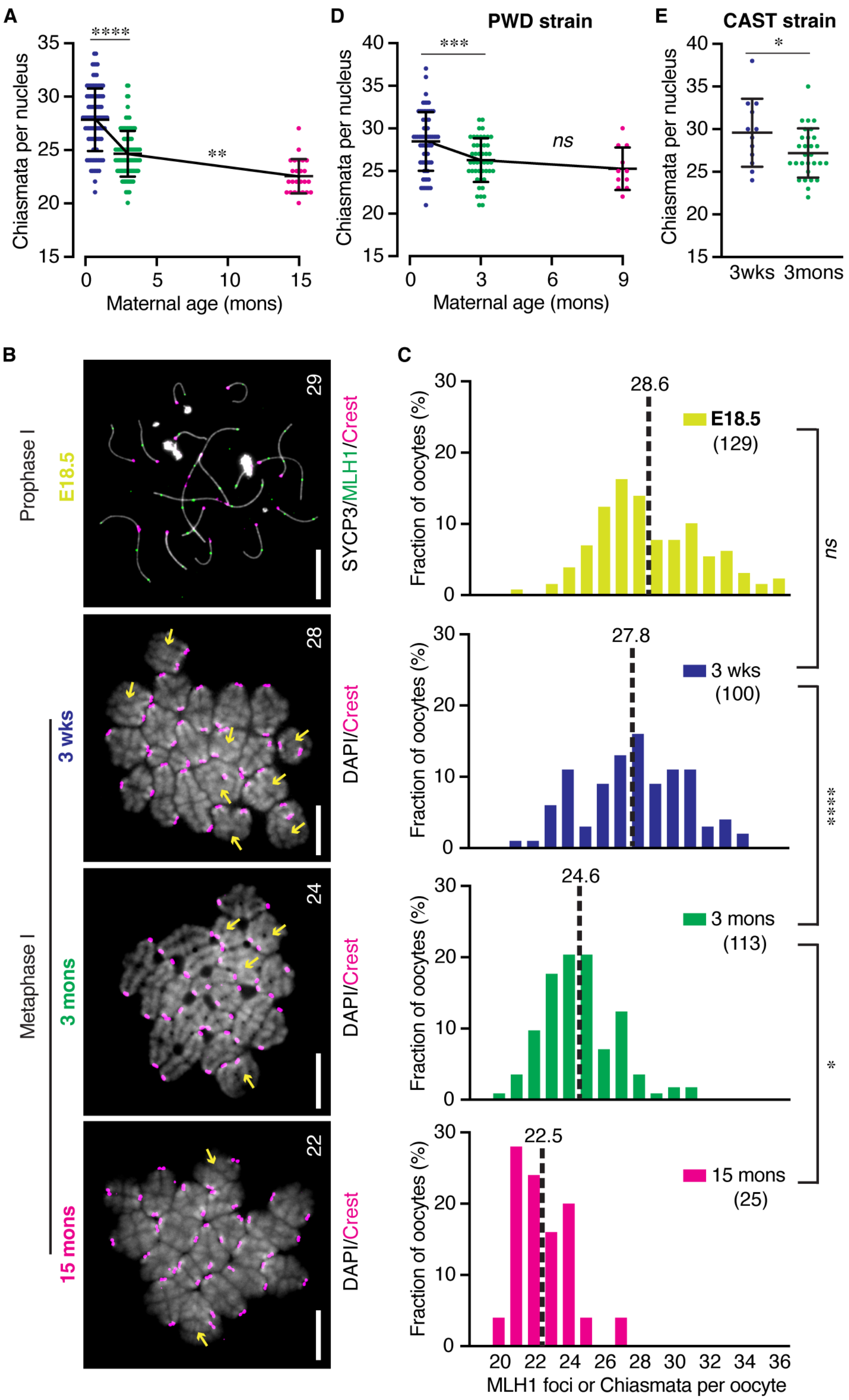


**Supplementary Figure S5 (related to Figure 3)**

(**A**). Chronological plots of chiasmata per oocyte quantified from metaphase-I oocytes. Error bars show means ± SD. **, *p*<0.01; ****, *p*<0.0001; Kruskal-Wallis test with Dunn's multiple comparisons

(**B**). Representative images of prophase I (top) and metaphase-I oocyte chromosome spreads to quantify crossover-specific MLH1 foci and chiasmata, respectively. Scale bars, 10 μm

(**C**). Distributions of the numbers of MLH1 or chiasmata per oocyte. Dashed black lines indicate the mean value of the population. *ns*, not significant; *, *p*<0.05; ****, *p*<0.0001; Kruskal-Wallis test with Dunn's multiple comparisons

(**D**). Chronological plots of chiasmata per oocyte quantified from metaphase-I oocytes from the PWD mouse strain. Error bars show means ± SD. *ns*, not significant; ***, *p*<0.001; Ordinary one-way ANOVA with Tukey's multiple comparisons test

(**E**). Chiasmata per oocyte quantified from metaphase-I oocytes from the CAST mouse strain. Error bars show means ± SD. *, *p*<0.05; Unpaired t test

### References


1. L. A. Bannister, L. G. Reinholdt, R. J. Munroe, J. C. Schimenti, Positional cloning and characterization of mouse mei8, a disrupted allele of the meiotic cohesin Rec8. *Genesis* **40**, 184–194 (2004).

2. J. Leem, T. Lemonnier, A. Khutsaidze, L. Tian, X. Xing, S. Bai, T. Nottoli, B. Mogessie, A versatile cohesion manipulation system probes female reproductive age-related egg aneuploidy. *Nat Aging* **5**, 2215–2227 (2025).

3. Y. Yun, M. Ito, S. Sandhu, N. Hunter, Cytological Monitoring of Meiotic Crossovers in Spermatocytes and Oocytes. *Methods Mol Biol* **2153**, 267–286 (2021).

4. A. Uchida, M. Yajima, An optogenetic approach to control protein localization during embryogenesis of the sea urchin. *Dev Biol* **441**, 19–30 (2018).

5. D. Clift, W. A. McEwan, L. I. Labzin, V. Konieczny, B. Mogessie, L. C. James, M. Schuh, A Method for the Acute and Rapid Degradation of Endogenous Proteins. *Cell* **171**, 1692-1706.e18 (2017).

6. T. Chiang, R. M. Schultz, M. A. Lampson, Age-dependent susceptibility of chromosome cohesion to premature separase activation in mouse oocytes. *Biol Reprod* **85**, 1279–1283 (2011).

7. S. Dunkley, B. Mogessie, Actin limits egg aneuploidies associated with female reproductive aging. *Sci Adv* **9**, eadc9161 (2023).

8. J. E. Holt, S. I. R. Lane, K. T. Jones, Time-lapse epifluorescence imaging of expressed cRNA to cyclin B1 for studying meiosis I in mouse oocytes. *Methods Mol Biol* **957**, 91–106 (2013).

9. B. Mogessie, Correction to: Visualization and Functional Analysis of Spindle Actin and Chromosome Segregation in Mammalian Oocytes. *Methods Mol Biol* **2101**, C1 (2020).

10. Y. Yun, S. Lee, C. So, R. Manhas, C. Kim, T. Wibowo, M. Hori, N. Hunter, Oocyte Development and Quality in Young and Old Mice following Exposure to Atrazine. *Environ Health Perspect* **130**, 117007 (2022).

11. J. R. Gruhn, A. P. Zielinska, V. Shukla, R. Blanshard, A. Capalbo, D. Cimadomo, D. Nikiforov, A. C.-H. Chan, L. J. Newnham, I. Vogel, C. Scarica, M. Krapchev, D. Taylor, S. G. Kristensen, J. Cheng, E. Ernst, A.-M. B. Bjørn, L. B. Colmorn, M. Blayney, K. Elder, J. Liss, G. Hartshorne, M. L. Grøndahl, L. Rienzi, F. Ubaldi, R. McCoy, K. Lukaszuk, C. Y. Andersen, M. Schuh, E. R. Hoffmann, Chromosome errors in human eggs shape natural fertility over reproductive life span. *Science* **365**, 1466–1469 (2019).

### Supplementary Movies S1 and S2

**Movie S1. Normal chromosome segregation at anaphase I in a mouse Y-oocyte**
*Time lapse movie of an oocyte from young adult female mice. Chromosomes (magenta) are labelled with H2B-mCherry. 00:00 indicates anaphase I onset.* Scale bar, 20 µm

**Movie S2. Chromosome lagging at anaphase I in a mouse J-oocyte**
*Time lapse movie of an oocyte from juvenile female mice. Chromosomes (magenta) are labelled with H2B-mCherry. 00:00 indicates anaphase I onset.* Scale bar, 20 µm
